# Cilia dysfunction in the lateral ventricles after neonatal intraventricular hemorrhage does not lead to functional changes in cilia-based CSF flow networks

**DOI:** 10.1101/2024.11.03.621724

**Authors:** Shelei Pan, Sruthi Ramagiri, Lihua Yang, David A. Giles, Isabella Xu, Maria Garcia Bonilla, Dakota DeFreitas, Lillian W. Siderowf, Grace L. Halupnik, Shriya Koneru, Gretchen M. Koller, Srinandan Polavarapu, Deepesh K. Gupta, Praveen Krishnamoorthy, Mark J. Miller, Prabagaran Esakky, David D. Limbrick, Phillip V. Bayly, Amjad Horani, Steven L. Brody, Moe R. Mahjoub, Jennifer M. Strahle

## Abstract

Intraventricular hemorrhage (IVH) has long been thought to lead to motile cilia dysfunction whereby intraventricular blood breakdown products damage and slough cilia from the ependymal wall. However, specifically how IVH may affect cilia development, structure, and transcriptional activation is not well-understood. Moreover, the impact of blood breakdown product-mediated cilia damage on the functional organization of cilia-based CSF flow networks is unknown. Here, we show hemoglobin exposure affects the number of ciliated ependymal cells in the lateral ventricle (LV) but does not impact *in vitro* beat frequency of the remaining cilia. Ultrastructurally, IVH decreases the total number of ciliary tufts without impacting axoneme structure. IVH does not result in changes in the expression of cilia-related genes and instead leads to downregulation of neurogenesis markers in parallel with innate immune upregulation. Functionally, we identify three previously uncharacterized cilia-mediated CSF flow domains in the LV lateral wall and show that IVH does not result in widespread disruption of their functional organization. These data de-emphasize cilia as a major contributor to global CSF dysfunction after IVH, and instead call attention to preserving the neurodevelopmental environment and preventing runaway innate immune system activation, as considerations to developing treatment strategies to prevent posthemorrhagic hydrocephalus and other neurodevelopmental sequelae.

## Introduction

Motile cilia are specialized hairlike organelles found in respiratory and oviduct epithelial cells, sperm, and the ependyma of the cerebral ventricles^1–3^. In each of these locations, motile cilia are thought to mediate the directional transport of fluids and solutes over the surface of the epithelium with which they are associated^1^. In the ependyma, which interfaces with the cerebrospinal fluid (CSF) of the cerebral ventricles, motile cilia are canonically thought to move CSF through the ventricular system to maintain CSF flow along the direction of their planar polarization^4–6^.

The hypothesized link between motile cilia and CSF transport through the ventricles is derived primarily from observations of patients and animal models with cilia dysfunction developing hydrocephalus, an abnormal buildup of CSF within the ventricles^7–15^. While hydrocephalus is associated with both congenital ciliopathies and has been postulated to be associated with acquired motile cilia loss mediated by blood breakdown exposure after intraventricular hemorrhage (IVH)^16,17^, there have been no mechanistic studies definitively linking motile cilia and hydrocephalus pathophysiology. It is unclear whether motile cilia dysfunction leads to aberrant CSF flow, and if aberrant CSF flow mediated by motile cilia loss may lead to altered CSF dynamics, CSF accumulation, and hydrocephalus.

In congenital hydrocephalus associated with ciliopathy, recent studies have suggested that the role of motile cilia in the pathophysiology of ventricular enlargement is in mediating impaired neurodevelopment secondary to disruption of cilia-mediated signaling, and not aberrant fluid movement^18–20^. How motile cilia are affected after neonatal IVH is less well-known, and it is unknown how exposure to intraventricular blood breakdown products impacts the severity, regional distribution, and physiological nature of damage to motile cilia.

In this study, we evaluated the effects of neonatal IVH on the structure and function of motile cilia in the lateral ventricles. We show that exposure to hemoglobin, a key pathogenic blood breakdown product released into the ventricles in the setting of IVH, results in a decrease in the number of ciliated ependymal cells. The axonemal structure and beat frequency of the remaining cilia were preserved, findings which were consistent with bead flow analyses demonstrating a lack of disruption to three previously uncharacterized motile cilia-mediated flow domains on the lateral wall of the lateral ventricle. Transcriptional analyses revealed no changes in the expression of motile cilia-related structural, functional, or ciliogenesis genes, but that innate immune signatures were upregulated in parallel with downregulation of key neurogenesis markers. We propose that neonatal IVH exerts its effects on motile cilia via hemoglobin-induced impaired ciliogenesis in conjunction with inflammation that results in cilia loss, and that this IVH-induced cilia dysfunction does not result in significant changes to motile cilia-mediated flow patterns that would be proportional to the degree of CSF dysfunction that occurs in posthemorrhagic hydrocephalus (PHH).

## Methods

### Animals

All animal procedures were approved by and performed in compliance with the Institutional Animal Care and Use Committee (IACUC) of Washington University in St. Louis School of Medicine protocols #22-0416 (rat) and #22-0254 (mouse). Lactating adult female Sprague Dawley rats with litters of 10 postnatal day 1-4 rat pups were purchased from Charles River Laboratories (strain 400, Charles River Laboratories, Wilmington, MA). All C57BL/6 mice were bred in-house and wild type. Animals were housed in a 12 h light/dark cycle in a temperature and humidity-controlled room. Water and food were provided ad libitum.

### Ependymal cell culture

Postnatal day 3 wild type C57BL/6 mice were sacrificed, and their brains harvested for lateral ventricle lateral wall wholemounts. Cells from the lateral walls were collected, dissociated, and plated into 25 cm^2^ flasks following the protocol described previously^21^. Briefly, cells were dissociated with an enzymatic digestion solution consisting of 17L*µ*L of Papain (#37A17241, Worthington, Lakewood, NJ, USA), 15L*µ*L of DNAse (#EN0521, Thermo Fisher Scientific, Waltham, MA, USA), and 288L*µ*g of l-Cysteine (#C7352, Sigma, St. Louis, MO, USA) in 1 mL of Dulbecco’s Modified Eagle Medium (DMEM)/Glutamax per brain for 1 hour at 37°C. Dissociated cells were plated into 25Lcm^2^ flasks and grown in proliferation media composed of DMEM/Glutamax supplemented with 10% fetal bovine serum and 1% penicillin/streptomycin (P/S) at 37°C. After confluence, cells were detached with 1 mL of trypsin-ethylenediaminetetraacetic acid (EDTA) and plated into 24-well plates (20L*µ*L of 2L×L106Lcells/mL) with coverslips for subsequent immunocytochemistry. The following day, DMEM/Glutamax with 1% P/S (differentiation media) was used for continued culture; cells progressively differentiated from neural stem cells to ependymal cells in this media. Cells were exposed to 100 μl of 100 *µ*M hemoglobin (#100714, MP Biomedicals, Solon, OH, USA) or aCSF (#3525, Tocris Bioscience, Bristol, UK) in 1 ml of differentiation media at day 3 of differentiation. Samples were analyzed after 7 days of exposure (day 10 of differentiation). The coverslips with attached cells were fixed with 4% paraformaldehyde for 10 min, washed 3 times with PBS for 5 minutes, and processed for immunocytochemistry with acetylated alpha-tubulin (5335S, Cell Signaling Technology, Danvers, MA) as previously described^21^.

### Cilia beat frequency analysis

For functional analysis of cilia movement *in vitro*, the beat frequency of cilia on cultured ependymal cells on Transwell membranes was imaged with a Nikon Ti inverted microscope using a 40x NAMC3 modulation contrast lens (Nikon, Tokyo, Japan) as previously described^22–24^. The environmental chamber was temperature, humidity, and CO_2_ controlled and held at 37° C during imaging. Images were recorded with a high-speed CMOS video camera from at least 5 fields of each Transwell insert and processed using Sisson-Ammons Video Analysis software (Ammons Engineering, Clio, MI).

### Intraventricular hemorrhage induction

IVH was induced through hemoglobin injection into the right lateral ventricle as previously described^25–27^. Briefly, bovine hemoglobin (#100714, MP Biomedicals, Solon, OH, USA) was constituted in aCSF (#3525, Tocris Bioscience, Bristol, UK) at a 150 mg/mL concentration and loaded into a 30 G, 0.33 cc insulin syringe attached to a micro-infusion pump. Postnatal day 2 (Figure 3) or 4 (all other figures) rats were anesthetized with isoflurane (3% induction, 2% maintenance) and secured using a stereotaxic frame. Bregma was exposed with a 1 cm midline scalp incision. 20 *µ*L of hemoglobin solution were injected into the right lateral ventricle at a 8 *µ*L/min rate using the following coordinates from bregma: 1.2-1.5 mm lateral, 0.5 mm anterior, 1.8-2.0 mm deep. Rats receiving an equivalent volume of intraventricular aCSF injection were used as volume controls. The needle was left in place for 5 minutes after the end of injection to prevent backflow before rats were sutured, allowed to recover from anesthesia, and returned to their cages.

### T2 structural MRI

72 hours after IVH or aCSF control induction, rats underwent T2-weighted structural MRI under isoflurane anesthesia (3% induction, 2-2.5% maintenance) to assess ventricle volume. Images were acquired using a Bruker 9.4-T (400-MHz), 20 cm clear bore MRI scanner (Bruker, Billerica, MA) with T2-weighted fast spin echo sequences with the following parameters: repetition time 4937.825 ms, echo time 66.00 ms, 2 averages, 1 repetition, echo spacing 16.500 ms, rare factor 8, field of view 15.000 x 15.000 mm, matrix 192 x 192, 32 axial slices, and 0.5 mm slice thickness.

### Histology

At postnatal day 7 (72 hours after P4 and 5 days after P2 aCSF or IVH-PHH induction), rats were sacrificed via isoflurane overdose and decapitation. Brains were harvested immediately and fixed in 4% paraformaldehyde (PFA) overnight. After fixation, brains were rinsed in 1x PBS 2x for 1 hour each, before dehydration in 30% EtOH for 30 minutes followed by 50% EtOH for 30 minutes and 70% EtOH overnight. After dehydration, brains were processed for formalin fixed paraffin embedding before sectioning at 8-10 *µ*m thickness in the coronal plane at the level of the lateral ventricles. Sections were dehydrated on a slide warmer at 40° C overnight followed by incubation at 60° C for 1 hour before deparaffinization in xylene for 20 minutes. Sections were rehydrated with descending grades of EtOH (100%, 95%, 70%, 50%, 30%) to D2O for 5 minutes each before heat-mediated antigen retrieval with 1x Diva Decloaker (DV2004, BioCare Medical, Pacheco, CA, USA). Sections were incubated in a blocking solution made from 95 mL 1x TBST, 5 mL normal goat serum (#S-10000, Vector Laboratories, Newark, CA), and 2.5 g bovine serum albumin (#A7888, Sigma-Aldrich, St. Louis, MO) for 1 hour before overnight incubation at 4°C in the following primary antibodies and dilutions: 1:200 Anti-alpha tubulin (#ab24610, Abcam, Cambridge, UK) and 1:100 Anti-Foxj1(#ab235445, Abcam, Cambridge, UK). After incubation with primary antibody, sections were washed with 1x PBS 6x for 5 minutes each, before incubation with the following secondary antibodies and dilutions: 1:2000 goat anti-Mouse 488 (#A28175, Invitrogen, Waltham, MA) and 1:2000 goat anti-Rabbit 594 (#A11012, Invitrogen, Waltham, MA) for 1 hour at room temperature. Sections were washed with 1x PBS 6x for 5 minutes each, then incubated with DAPI diluted 1:500 in PBS, before they were mounted with Prolong Gold Antifade (#P36970, Molecular Probes, Eugene, OR, USA) and coverslipped for confocal imaging.

### Scanning electron microscopy

For SEM, rats were sacrificed 72 hrs or 11 days post-P4 aCSF control or IVH-PHH induction. Brains were harvested immediately and fixed in a 2.5% glutaraldehyde, 2% PFA, and 0.15M cacodylate buffer (16537-15, Electron Microscopy Sciences, Hatfield, PA, USA) overnight at 4° C. Samples were rinsed in 0.15 M cacodylate buffer 3x for 20 minutes each before incubation in a 2% OsO4 in 0.15 M cacodylate buffer for 1.5 hours in the dark (19152, Electron Microscopy Sciences, Hatfield, PA, USA). Post-OsO4 incubation, samples were rinsed 4x in ultrapure water for 15 minutes each before dehydration by immersion in a series of increasing ethanol grades (30%, 50%, 70%, 95%, 100%) for 10 minutes each. Once dehydrated, samples were then loaded into a critical point drier (Leica EM CPD300, Vienna, Austria) for 12 CO2 exchanges at the lowest speed setting. Once dry, samples were mounted on aluminum stubs with carbon adhesive tabs and coated with 12 nm of carbon and 12 nm of iridium (Leica ACE 600, Vienna, Austria). SEM images were acquired on a FE-SEM (Zeiss Merlin, Oberkochen, Germany) at 1.5 kV and 0.1 nA.

### Transmission electron microscopy

72 hrs post-P4 aCSF control or IVH-PHH induction, rats were sacrificed and their brains harvested immediately and fixed in a 2.5% glutaraldehyde, 2% paraformaldehyde, and 0.15M cacodylate buffer containing 2 mM calcium chloride (16537-15, Electron Microscopy Sciences, Hatfield, PA, USA) overnight at 4° C. Post-fixation, 100 *µ*m coronal sections were cut at the level of the lateral ventricle using a vibratome (Leica VT1200S, Vienna, Austria), then rinsed in a 0.15 M cacodylate buffer containing 2 mM calcium chloride 3x for 10 minutes each. Next, samples were incubated in a 2% OsO4 in 0.15 M cacodylate buffer solution containing 2 mM calcium chloride (19152, Electron Microscopy Sciences, Hatfield, PA, USA) for 2 hours in the dark, before being rinse 3x in ultrapure water for 10 minutes each. The samples were stained en bloc with 2% aqueous uranyl acetate overnight at 4° C in the dark (22400-2, Electron Microscopy Sciences, Hatfield, PA, USA). After staining, samples were washed 4x in ultrapure water and dehydrated by immersion in a series of increasing ethanol grades (30% x1, 50% x1, 70% x1, 95% x1, 100% x3) for 10 minutes each. Once dehydrated, samples were infiltrated with resin (LX112, Electron Microscopy Sciences, Hatfield, PA, USA). The samples were flat embedded and polymerized for 48 hours at 60° C. Post curing, a region of interest along the ependymal wall of the lateral ventricle was excised and mounted perpendicularly on a blank epoxy stub for sectioning. 70 nm ultrathin sections were taken and post-stained with 2% aqueous uranyl acetate and lead citrate (22400, 22405, 17800, 17900, 21140, Electron Microscopy Sciences, Hatfield, PA, USA). Samples were imaged on a TEM (Jeol JEM-1400 Plus) at 1200 kV.

### Bulk RNA sequencing tissue processing and computational analysis

72 hours after aCSF control or IVH-PHH induction, rats were sacrificed, and the right lateral wall of the lateral ventricle was microdissected for bulk RNA sequencing. Total RNA was isolated, and samples with a Bioanalyzer RIN score greater than 8.0 were selected for subsequent analysis (3 per group). Ribosomal RNA was removed by poly-A selection using Oligo-dT beads (mRNA Direct kit, Life Technologies). mRNA was then fragmented in reverse transcriptase buffer and heating to 94 degrees for 8 minutes. mRNA was reverse transcribed to yield cDNA using SuperScript III RT enzyme (Life Technologies, per manufacturer’s instructions) and random hexamers. A second strand reaction was performed to yield ds-cDNA. cDNA was blunt ended, had an A base added to the 3’ ends, and then had Illumina sequencing adapters ligated to the ends. Ligated fragments were then amplified for 12-15 cycles using primers incorporating unique dual index tags. Fragments were sequenced on an Illumina NovaSeq-6000 using paired end reads extending 150 bases. RNA-seq reads were then aligned to the Ensembl Rat Rnor_6.0 genome and quantitated with an Illumina DRAGEN Bio-IT on-premise server running version 3.9.3-8 software. RNA-seq reads were analyzed with R version 4.3.0 and RStudio version 2023.03. Reads were imported with tximport^28^, and differential gene expression determined by DESeq2^29^. Significance was defined by an adjusted p-value using the Benjamini-Hochberg method for multiple comparisons. Results were visualized with EnhancedVolcano^30^ and ComplexHeatmap^31^.

### Lateral wall wholemount and bead flow image acquisition

10-11 days post-aCSF control and IVH-PHH induction, rats were sacrificed and the lateral wall of the lateral ventricle was isolated as a wholemount as previously described^32^. The wholemount was secured to a 35 mm petri dish with cyanoacrylate glue (#B00016067, 3M, St. Paul, MN), with the cortex side glued to the petri dish and the ciliated ependymal surface facing up. The petri dish was filled with PBS to prevent tissue dehydration and placed on the motorized stage of a Zeiss Axio Zoom v16 fluorescent microscope (Zeiss, Oberkochen, Germany) set to 7x zoom. Image acquisition was started using a Hamamatsu Orca Flash sCMOS high-speed fluorescence camera, and 2 *µ*L of a solution of 10 *µ*m red fluorescent (580/605) polystyrene microspheres (#F8834, Invitrogen, Waltham, MA, USA) and 4.0 *µ*m green fluorescent sulfate-modified microspheres (#8859, Invitrogen, Waltham, MA, USA) diluted 1:2500 in aCSF was dropped onto the ependymal surface using a 2 *µ*L pipette. Images were acquired with an interval time of 1.0s and exposure time 50 ms for 500 frames across Cy3 and EGFP channels. The last 250 frames were used for analysis.

### Bead flow analysis

Raw images were imported into Imaris v.10.x and a new surface was created to restrict the working space to the lateral ventricle ependymal surface. For ROI analyses, surfaces were created to include only the ROI, generating three surfaces (rostral, caudal, superior) per animal. The surface was duplicated over all timepoints, and two masks were generated for each surface, one for the 4 *µ*m green fluorescent sulfate-modified microspheres and a second for the 10 *µ*m red fluorescent polystyrene microspheres. Spot creation parameters were created for the 4 *µ*m green fluorescent sulfate-modified microspheres (estimated diameter = 25.0 *µ*m, track speed mean above 1.00 *µ*m/s) and the 10 *µ*m red fluorescent polystyrene microspheres (estimated diameter = 50.0 *µ*m, track speed mean above 1.00 *µ*m/s) and applied to the surfaces to generate spots for each bead and their corresponding track. To export temporal data for statistical analyses, the time view function within vantage function was used to set the y-axis to the desired output statistics (speed, acceleration), before export of detailed statistics to .csv format. To export track/ROI track data, track length, straightness, and track speed variation output statistics were exported via the statistics tab to .csv format. Individual values across all times were averaged over all tracks/beads for each animal (n=3 aCSF control, n=5 IVH-PHH) to obtain one average value per animal for each output statistic of interest for statistical analysis.

### Statistical analysis

Statistical methods were not used to recalculate or predetermine sample sizes. All graphs shown in the results were expressed as mean ± SEM from 3-5 independent experiment replicates. Unless otherwise specified, associations between two continuous variables were assessed using an unpaired T-test, and associations between more than two continuous variables were assessed using a one-way ANOVA with post-hoc Tukey. All tests were 2-tailed, and p-values of less than 0.05 were considered statistically significant. All analyses were performed using Microsoft Office Excel (Version 16.36) or GraphPad Prism (Version 9.0.0) unless otherwise specified.

## Results

### Hemoglobin exposure *in vitro* leads to a reduction in both the total number of cells in a multiciliated state and the number of cilia per cell

We used a previously reported ependymal cell culture model in which cells differentiate from monociliated neural stem cells to multiciliated ependymal cells in a time-dependent fashion, where approximately 4% of all cells differentiate into the multiciliated ependymal phenotype by 3 days of differentiation, 23% at 5 days, and 52% at 7 days^21^. Therefore, to evaluate the effect of hemoglobin, a key pathogenic blood breakdown product released into the ventricles in IVH, on ependymal cell development, ciliogenesis, and cilia loss *in vitro*^21^, we chose to expose cells to standard media, aCSF, or 100 *µ*m hemoglobin early on in the process of differentiation at day 3. At day 10 of differentiation, cells were immunostained for the microtubule marker acetylated alpha tubulin to label cilia and allow for identification of multiciliated and uniciliated cells (Figure 1A).

**Figure 1.**
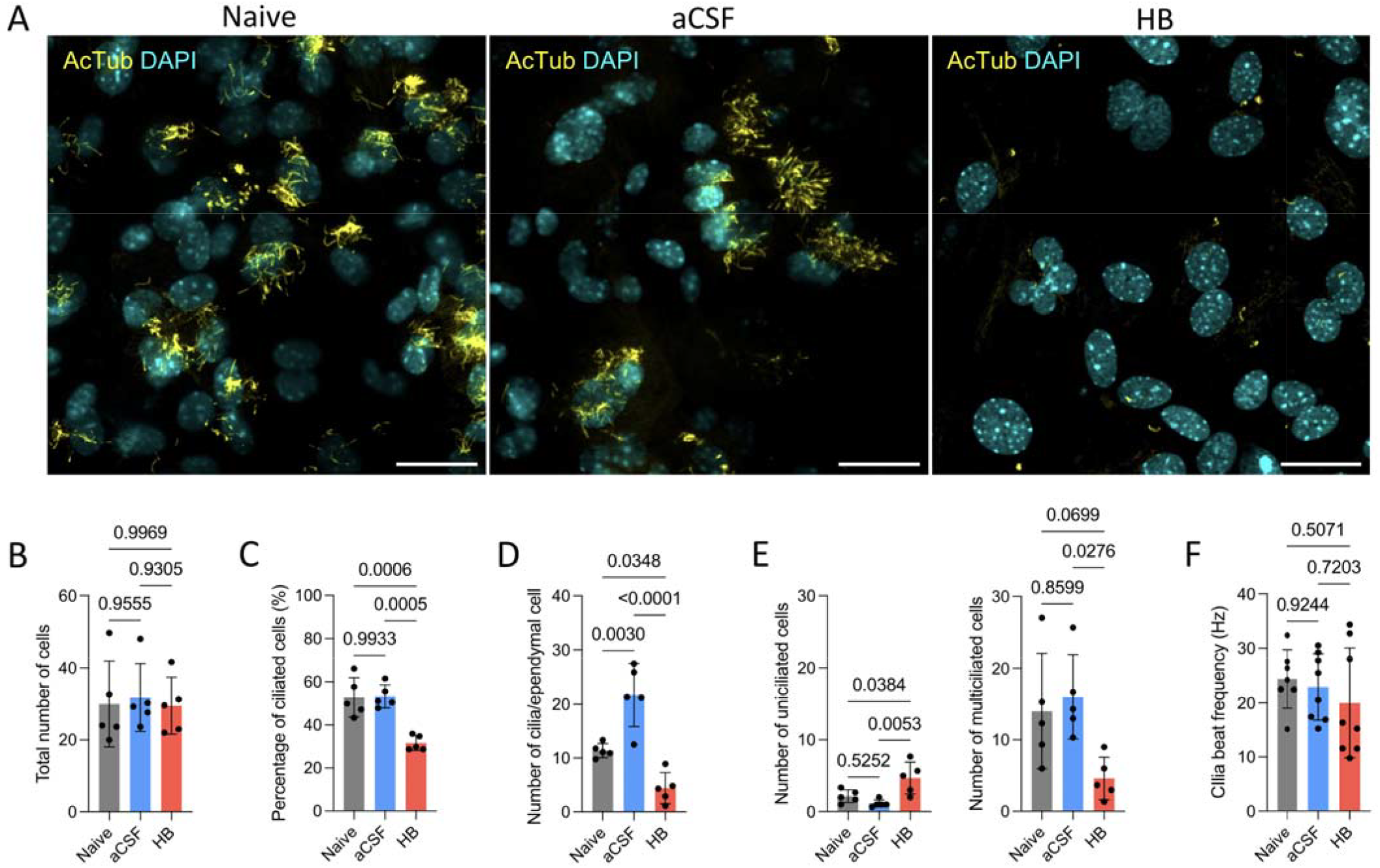
Hemoglobin exposure *in vitro* affects ependymal cell differentiation and ciliogenesis, but not beat frequency. **A**, Representative photomicrographs of cilia in day 10 post-differentiation rat ependymal cells 7 days after exposure to standard media (naïve), artificial CSF (aCSF), or 100 *µ*M hemoglobin (HB) at day 3 of differentiation. Scalebars = 25 *µ*m. **B**, There was no significant difference in the total number of cells in the culture wells of naïve, aCSF, or HB-treated conditions. **C**, The percent of ciliated cells, calculated as the number of cells with cilia/the number of total cells, was smaller after hemoglobin exposure compared to the naïve and aCSF conditions. There was no difference between the naïve and aCSF conditions. **D**, The number of cilia per ependymal cell was significantly fewer after hemoglobin exposure compared to both the naïve and aCSF conditions, while the number of cilia per ependymal cell was greater after aCSF exposure compared to the naïve condition. **E**, The number of cells which were uniciliated (left) was significantly greater after HB exposure compared to the naïve and aCSF conditions, while the number of cells which were multiciliated (right) was significantly fewer after HB exposure compared to the aCSF condition. **F**, There were no significant differences in cilia beat frequency between naïve, aCSF, and HB conditions. For all graphs in B-F, data were analyzed via one-way ANOVAs with post-hoc Tukeys. Data are shown as mean ± S.D., with cells obtained from at least n = 3 animals each for the naïve, aCSF, and HB conditions.

We found that while the total number of cells remained constant across all conditions (Figure 1B), hemoglobin exposure resulted in a lower percentage of ciliated cells compared to standard media (p=0.0006) and aCSF (p=0.0005) (Figure 1C). The number of cilia per ependymal cell was also fewer in cells exposed to hemoglobin vs. aCSF (p<0.0001, Figure 1D). We also quantified the total number of uniciliated and multiciliated cells, and found that hemoglobin exposure led to a greater number of uniciliated cells and fewer multiciliated cells (Figure 1E), potentially reflecting a less differentiated state. In order to determine whether hemoglobin also affects the coordinated beating function of motile cilia, we performed beat frequency analyses and found no significant differences in the beat frequency of motile cilia exposed to hemoglobin vs. aCSF and standard media (Figure 1F), suggesting the function of surviving cilia in ependymal cells which successfully underwent differentiation despite hemoglobin exposure is not altered.

### IVH-PHH results in widespread decreases in cilia in the lateral ventricle, with variable effects on ependymal Foxj1 expression by region and time of hemorrhage

We next aimed to evaluate the effect of intraventricular hemoglobin on cilia *in vivo* using our previously described animal model of IVH-PHH^25–27^. It has previously been reported that the number of multiciliated ependymal cells in the lateral wall of the lateral ventricle starts to increase rapidly between postnatal days 0 and 2 with continued rapid differentiation through postnatal day 4, after which they become widely distributed to cover the entire lateral wall^33^. Postnatal day 4 also coincides approximately with localization of Foxj1 expression to ventricular epithelial cells undergoing apical surface expansion as a hallmark of E1 ependymal cell differentiation^34^, while ventricular epithelial cells expressing Foxj1 at postnatal day 2 remain in a less differentiated state.

We induced IVH at postnatal days 2 and 4 to investigate the effect of hemorrhage at two timepoints during the peak of differentiation: an earlier timepoint when fewer total ependymal cells are differentiated and the wall consists of a mix of immature and mature ependymal cells, and a later timepoint when there is a greater number of mature ependymal cells. By postnatal day 7, there were widespread decreases in the percent of ependymal cells with cilia in the lateral wall (Figure 2C), medial wall (Figure 2F), and superior wall (Figure 2I) of the lateral ventricle in response to postnatal day 4 IVH, but decreases in only the lateral (Figure 3C) and medial (Figure 3F) walls in response to postnatal day 2 IVH. When the percent of ependymal cells with cilia were compared between each of the walls, we found there were significantly fewer ciliated ependymal cells on the lateral wall vs. medial wall after postnatal day 4 IVH, suggesting IVH-PHH differentially affected the lateral wall (Figure 2L, 2M). In contrast, there were no significant differences in response to postnatal day 2 IVH, suggesting IVH at this earlier timepoint does not result in differential damage to cilia by wall (Figure 3L). Of note, while there were no differences in Foxj1 expression after postnatal day 2 IVH (Figure 3), postnatal day 4 IVH led to a decrease in the percent of Foxj1+ ependymal cells on the lateral wall of the lateral ventricle compared to the controls by postnatal day 7 (Figure 2B).

**Figure 2.**
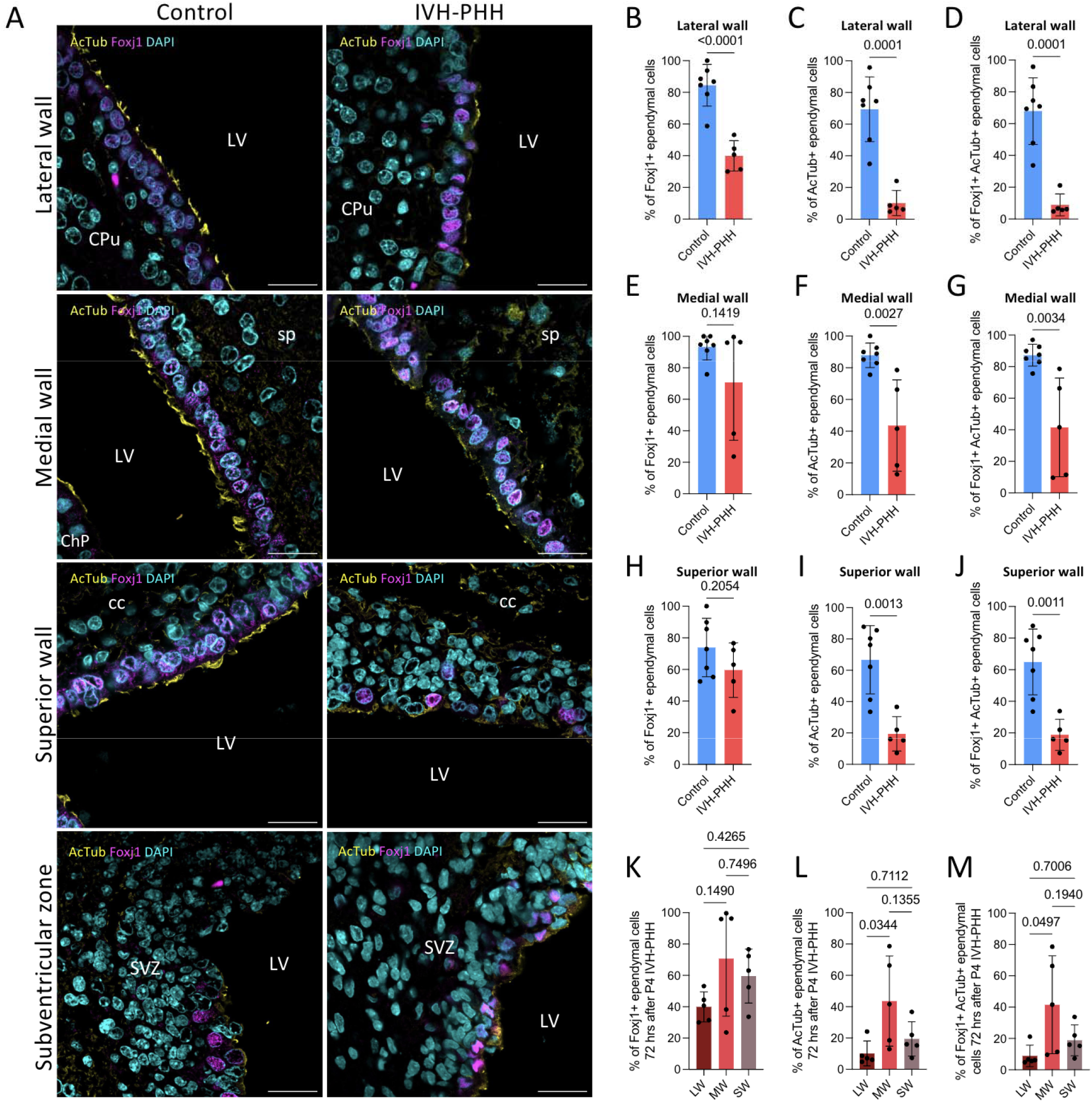
Intraventricular hemorrhage leads to a decrease in Foxj1 expression isolated to the lateral wall ependyma of the lateral ventricle, with widespread decreases in the number of ependymal cells with cilia. **A**, Representative images of acetylated tubulin (AcTub), and Foxj1 expression along the ependyma lining the lateral, medial, and superior walls and subventricular zone of the lateral ventricle 3 days after aCSF or HB injection into the right lateral ventricle of postnatal day 4 rats to induce the control and intraventricular hemorrhage-posthemorrhagic hydrocephalus (IVH-PHH) conditions, respectively. There was regional heterogeneity in Foxj1 expression, with multiple strata/pseudostrata of Foxj1+ cells in the lateral wall, but only single layers of Foxj1+ cells along the three other regions. Similarly, there was robust AcTub staining indicating the presence of cilia along the ependyma of the lateral, medial, and superior walls of the lateral ventricle in control rodents, but relatively less staining in the ependyma lining the subventricular zone. Scalebars = 25 *µ*m. Abbreviations: CPu, caudate putamen; LV, lateral ventricle; ChP, choroid plexus; sp, septum pellucidum; cc, corpus callosum; SVZ, subventricular zone. **B-D**, Quantification of the percent of ependymal cells lining the lateral wall of the lateral ventricle which express Foxj1 (B), AcTub (C), and both Foxj1 and AcTub (D) 3 days after aCSF control or IVH-PHH induction. **E-G**, Quantification of the percent of ependymal cells lining the medial wall of the lateral ventricle which express Foxj1 (E), AcTub (F), and both Foxj1 and AcTub (G). **H-J**, Quantification of the percent of ependymal cells lining the superior wall of the lateral ventricle which express Foxj1 (H), AcTub (I), and both Foxj1 and AcTub (J). **K-M**, Quantification of differential ependymal expression of Foxj1 (K), AcTub (L), and Fox1 and AcTub (M) across the lateral (LW), medial (MW), and superior (SW) walls of the lateral ventricles after IVH-PHH induced at postnatal day 4. While IVH did not differentially affect Foxj1 expression by region, AcTub expression (L) and Foxj1/AcTub co-expression (M) in the lateral wall were more severely affected compared to the medial wall. Data in B-J were analyzed via unpaired, two-tailed t-tests, while data in K-M were analyzed via One-way ANOVAs with post-hoc Tukeys All data are shown as mean ± S.D., with at least n = 4 animals each for the control and IVH-PHH conditions.

**Figure 3.**
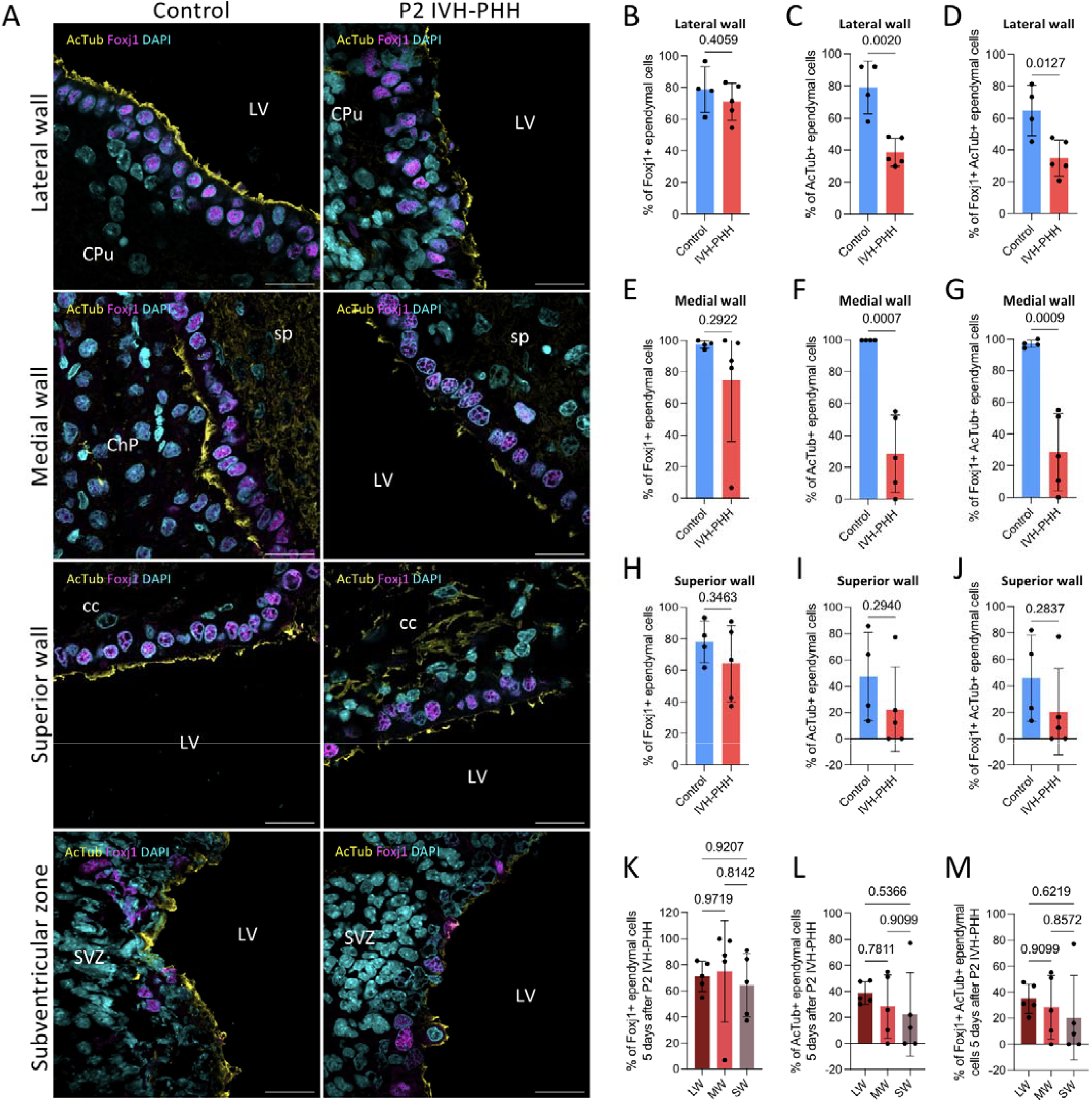
Early intraventricular hemorrhage induced at postnatal day 2 leads to a decrease in the number of ependymal cells with cilia, but no changes in Foxj1 expression. **A**, Representative images of acetylated tubulin (AcTub), and Foxj1 expression along the ependyma lining the lateral, medial, and superior walls and subventricular zone of the lateral ventricle 5 days after aCSF or HB injection into the right lateral ventricle of postnatal day 2 rats to induce the control and intraventricular hemorrhage-posthemorrhagic hydrocephalus (IVH-PHH) conditions, respectively. Scalebars = 25 *µ*m. Abbreviations: CPu, caudate putamen; LV, lateral ventricle; ChP, choroid plexus; sp, septum pellucidum; cc, corpus callosum; SVZ, subventricular zone. **B-D**, Quantification of the percent of ependymal cells lining the lateral wall of the lateral ventricle which express Foxj1 (B), AcTub (C), and both Foxj1 and AcTub (D) 5 days after aCSF control or IVH-PHH induction. **E-G**, Quantification of the percent of ependymal cells lining the medial wall of the lateral ventricle which express Foxj1 (E), AcTub (F), and both Foxj1 and AcTub (G). **H-J**, Quantification of the percent of ependymal cells lining the superior wall of the lateral ventricle which express Foxj1 (H), AcTub (I), and both Foxj1 and AcTub (J). **K-M**, Quantification of differential ependymal expression of Foxj1 (K), AcTub (L), and Fox1 and AcTub (M) across the lateral (LW), medial (MW), and superior (SW) walls of the lateral ventricles after IVH-PHH induced at postnatal day 2. There were no differences in regional expression of either marker. Data in B-J were analyzed via unpaired, two-tailed t-tests, while data in K-M were analyzed via One-way ANOVAs with post-hoc Tukeys All data are shown as mean ± S.D., with at least n = 4 animals each for the control and IVH-PHH conditions.

We next used SEM to investigate the effect of IVH on cilia across the entire lateral wall of the lateral ventricle (Figure 4). We found that while there were no differences in the number of ciliary tufts in rostral (Figure 4D), middle (Figure 4E), or caudal (Figure 4F) regions of the lateral wall 72 hours after IVH, there were significantly fewer tufts in the rostral (Figure 4H) and middle (Figure 4I) regions by 11 days post-IVH. There were no significant differences at either timepoint when we compared the number of ciliary tufts between the rostral, and middle, and caudal regions (Figure 4K, 4L), suggesting IVH did not result in differential cilia loss by region of the lateral wall.

**Figure 4.**
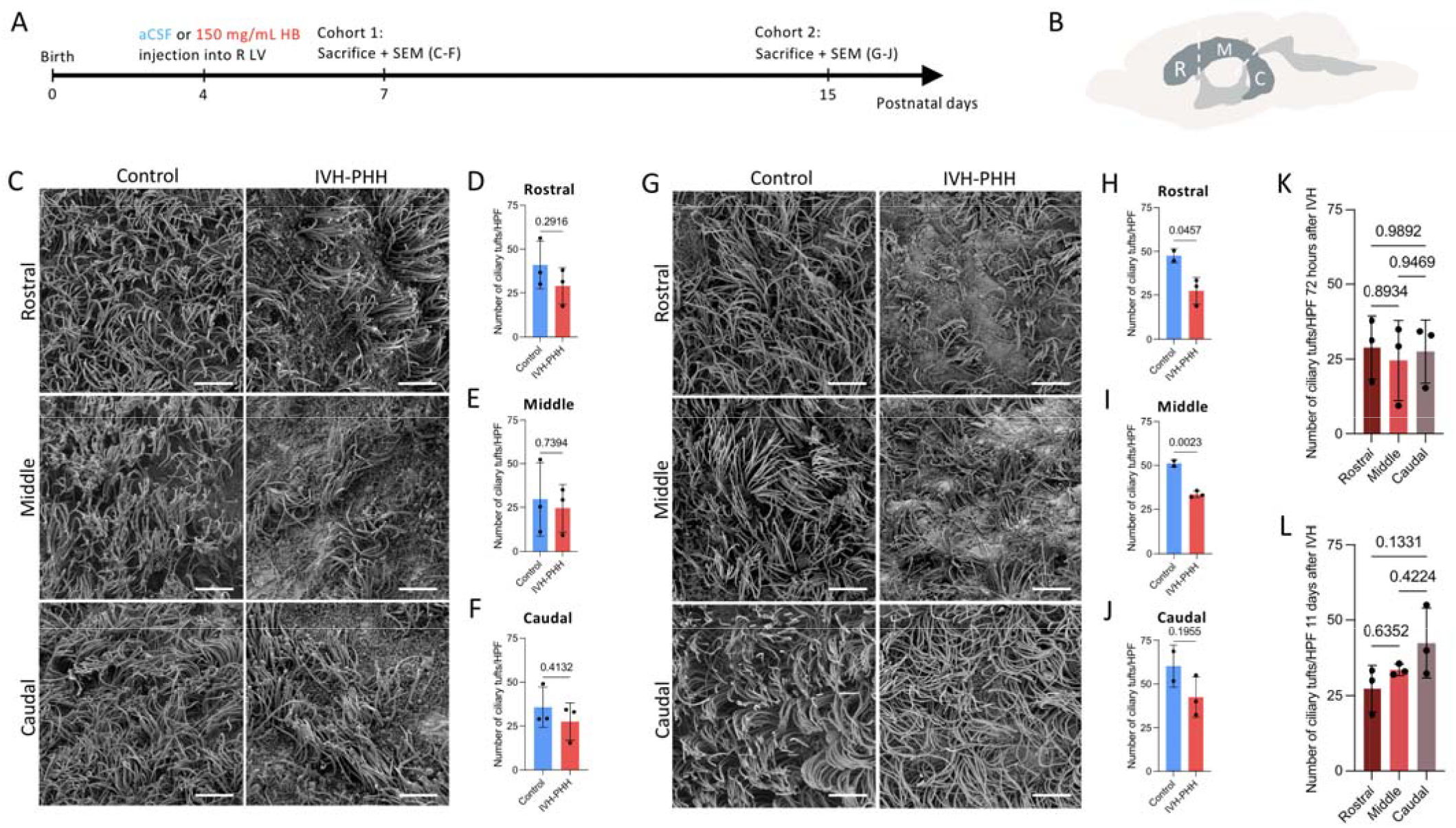
The number of ciliary tufts on the lateral ventricle wall is decreased at long-term, but not acute, timepoints after intraventricular hemorrhage. **A**, Timeline of artificial CSF (aCSF) or 150 mg/mL hemoglobin (HB) injection into the right lateral ventricle (R LV) of postnatal day 4 rats to induce the control or intraventricular hemorrhage-posthemorrhagic hydrocephalus (IVH-PHH) conditions. One cohort of rats was sacked 72 hours post-injection at postnatal day 7 and brains collected for processing for scanning electron microscopy (SEM) (data in subfigures C-F), while a second cohort of rats was sacked 11 days post-injection at postnatal day 15 (subfigures G-J). **B**, Schematic of the ventricular system including the lateral ventricle (dark gray) showing rostral (R), middle (M), and caudal (C) regions of the lateral wall of the lateral ventricle from which data was obtained for subfigures C-L. **C**, Representative SEM images of cilia in rostral, middle, and caudal regions of the lateral wall of the lateral ventricle 72 hours post-control and IVH-PHH induction. Scalebars = 5 *µ*m. **D-F**, Quantification of the number of ciliary tufts per 2,000x high power field in the rostral (D), middle (E), and caudal (F) regions of the lateral wall of the lateral ventricle 72 hours post-control and IVH-PHH induction. Data were analyzed via unpaired, two-tailed t-tests and are shown as mean ± S.D., with at least n = 3 animals each for the control and IVH-PHH conditions. **G**, Representative SEM images of cilia in rostral, middle, and caudal regions of the lateral wall of the lateral ventricle 11 days post-control and IVH-PHH induction. Scalebars = 5 *µ*m. **H-J**, Quantification of the number of ciliary tufts per 2,000x high power field in the rostral (H), middle (I), and caudal (J) regions of the lateral wall of the lateral ventricle 11 days post-control and IVH-PHH induction. Data were analyzed via unpaired, two-tailed t-tests and are shown as mean ± S.D., with at least n = 3 animals each for the control and IVH-PHH conditions. **K-L**, Quantification of the number of ciliary tufts per 2000x high power field 72 hours (K) and 11 days (L) post-IVH-PHH induction. IVH did not result in differential loss of cilia in any of the three regions at either timepoint. Data were analyzed via one-way ANOVAs with post-hoc Tukeys and are shown as mean ± S.D., with at least n = 3 animals each for the control and IVH-PHH conditions.

### Motile cilia axonemal structure is preserved after IVH-PHH

Given IVH occurs around the time of ependymal differentiation and ciliogenesis and that there is a relative lack of information surrounding the effect of IVH on the axoneme, we performed TEM to assess motile cilia ultrastructure in cross section. We identified motile cilia by their 9+2 microtubule arrangement structure with nine radial doublet microtubules and a central pair of singlet microtubules (Figure 5). We identified normal radial spoke, inner dynein arm, and outer dynein arm structure in all motile cilia imaged, suggesting IVH does not affect the organization of the motile cilia axoneme (Figure 5).

**Figure 5.**
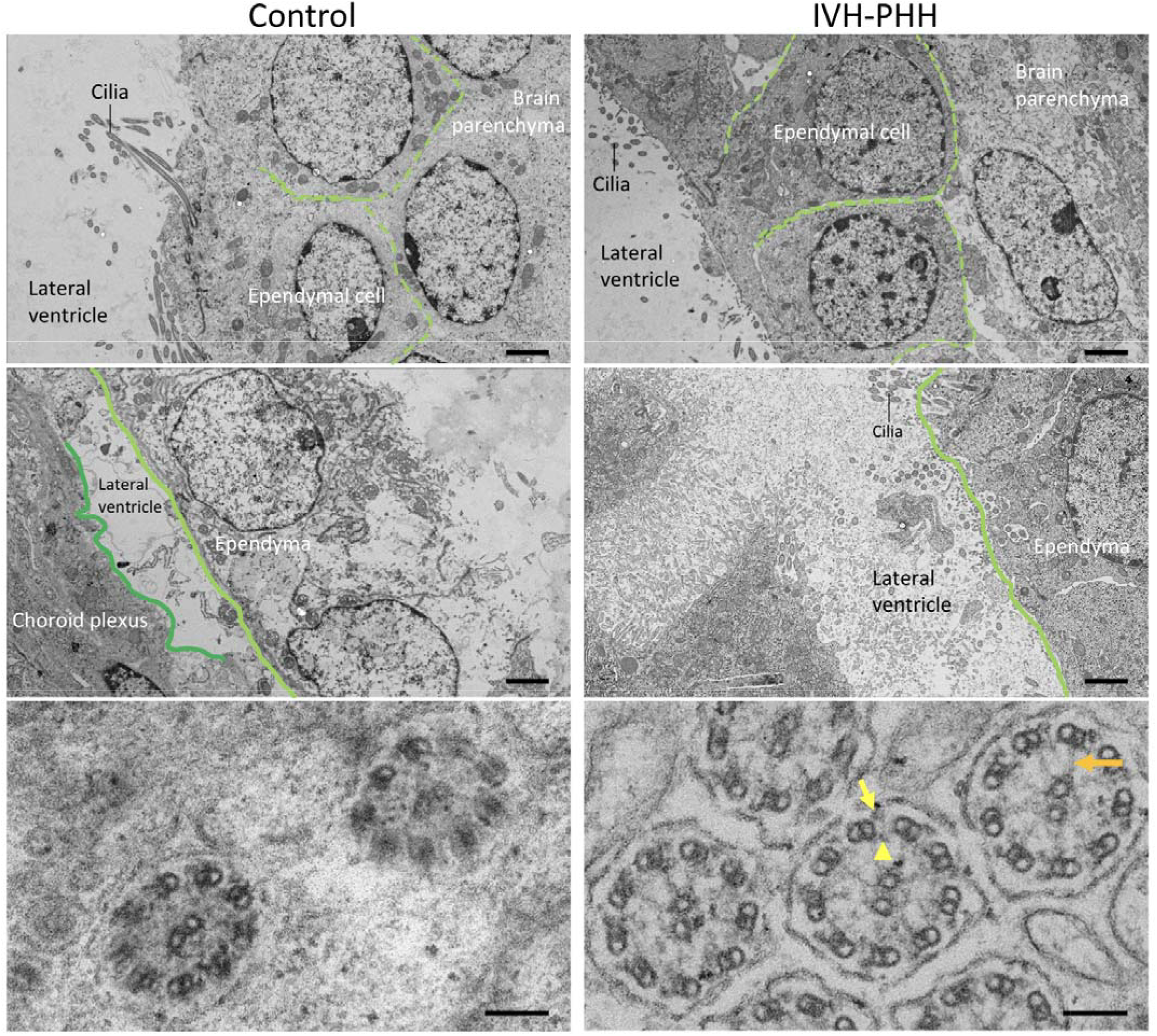
Transmission electron microscopy (TEM) reveals there are no ultrastructural changes in the ependyma and cilia of the lateral ventricle 72 hours after intraventricular hemorrhage. Representative images of ependyma (top row), ependyma-choroid plexus interactions (middle row), and cilia obtained 72 hours after artificial CSF and HB injection into the right lateral ventricle of P4 rats to induce the control and intraventricular hemorrhage-posthemorrhagic hydrocephalus (IVH/PHH) conditions, respectively. Light green dashed lines indicate individual ependymal cells, light green solid lines indicate the apical surface of the ependyma facing the lateral ventricle, and dark green solid lines delineate the choroid plexus. The presence of the outer dynein arm is indicated with a yellow arrow, while the inner dynein arm is indicated with a yellow arrowhead. Radial spokes are indicated with an orange arrow. Top and middle row scalebars = 2 *µ*m, bottom row scalebars = 100 nm.

### IVH-PHH does not affect cilia gene expression but leads to innate immune upregulation and neurogenesis downregulation

As IVH-PHH leads to decreases in cilia on ependymal cells both *in vitro* and *in vivo*, we questioned if the decrease in cilia may be due to 1) altered ependymal cell differentiation and ciliogenesis preventing cilia development, or 2) post-ciliogenesis cilia loss secondary to the toxic effects of CSF blood breakdown product exposure. We hypothesized that IVH-induced impaired ciliogenesis would be reflected in changes in the expression of key cilia assembly, regulatory, or differentiation genes, while blood breakdown product-induced ependymal injury leading to remodeling and/or mechanical cilia sloughing would be reflected in upregulation of immune and/or inflammatory markers.

We isolated the lateral wall of the lateral ventricle 72 hours after aCSF control and IVH-PHH induction and performed bulk RNA sequencing on the tissues. We detected no significant changes in the expression of key genes involved in motile ciliogenesis and ependymal differentiation (*Foxj1, Rfx3*), dynein outer arm assembly (*Ccdc103, Lrrc6*) and composition (*Dnaic1, Dnaic2, Dnah5*), and radial spoke head structure (*Rsph4a*) after IVH-PHH vs. aCSF controls (Figure 6C). However, there was widespread innate immune upregulation after IVH-PHH in the following genes: *Adgre1* (p<0.001, Figure 6I), *CD4* (p=0.006, Figure 6J), *CD45* (p=0.013, Figure 6K), *CD74* (p<0.001, Figure 6L), *Clec4a1* (p=0.0009, Figure 6M), *Cxcl13* (p<0.001, Figure 6N), *Cybb* (p<0.001, Figure 6O), *C3* (p<0.001, Figure 6P), *Il1b* (p<0.001, Figure 6Q), *Lyz2* (p<0.001, Figure 6R), *Mrc1* (p=0.002, Figure 6S), *Siglec1* (p=0.002, Figure 6T), *Tlr8* (p=0.031, Figure 6U), and *Tlr9* (p=0.003, Figure 6V).

**Figure 6.**
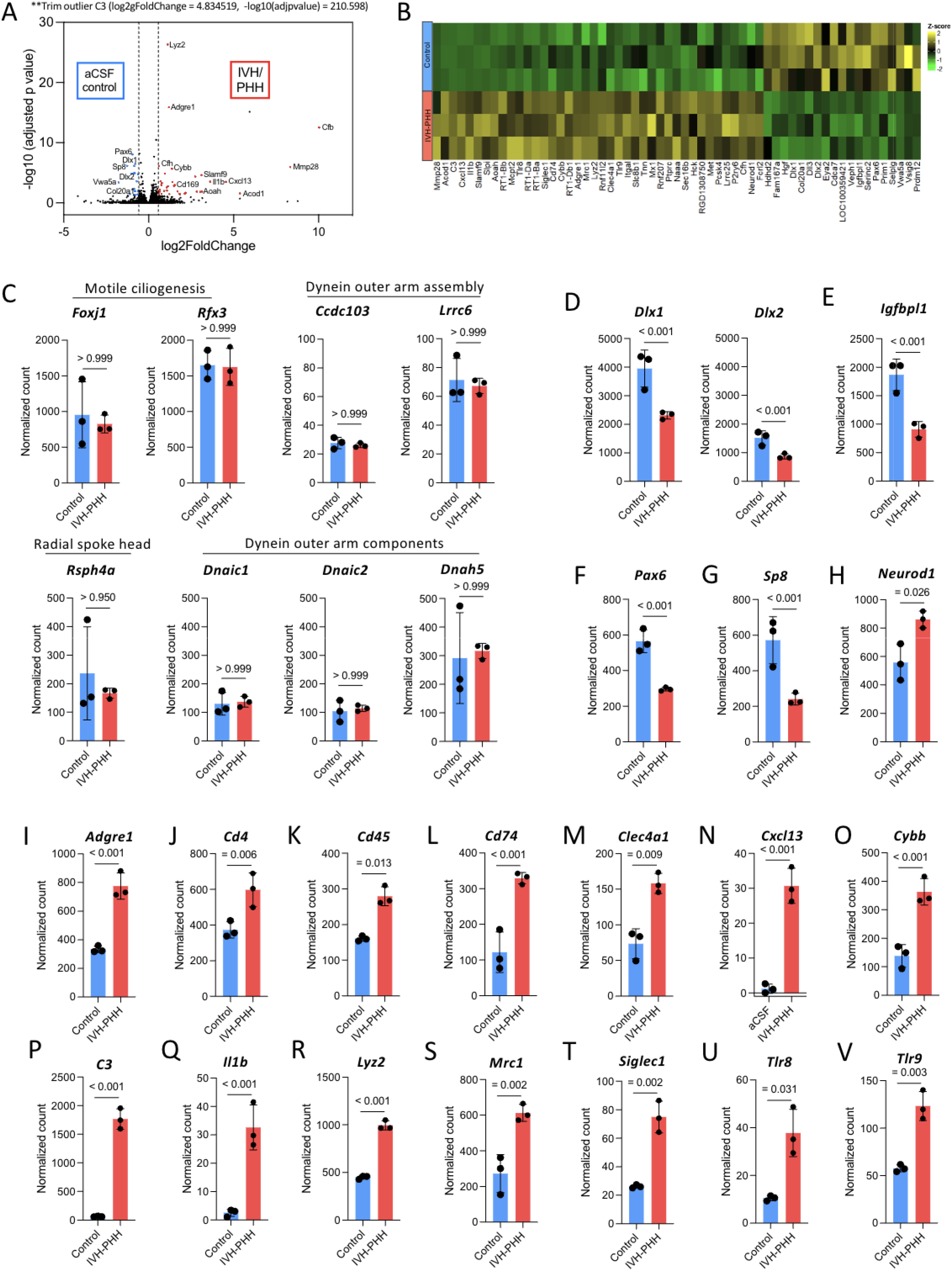
Neurogenesis markers are downregulated in parallel with innate immune upregulation 72 hours post-intraventricular hemorrhage, with no changes in the expression of cilia-related genes. **A**, Volcano plot showing adjusted p value vs. log2fold change for genes expressed in the aCSF control neonatal rat brain and 72 hours after IVH/PHH. Red and blue datapoints represent adjusted p values < 0.05. Select genes which are differentially expressed in either aCSF control (blue) or IVH/PHH (red) are identified on the graph. **B**, Heat map of normalized count values for genes that are differentially expressed (p-value<0.05, mean count >5, log2FC>0.585) in the neonatal rat brain 72 hours after IVH/PHH. **C**, Graphical representations of normalized counts for select cilia-related genes involved in motile ciliogenesis, dynein outer arm assembly, radial spoke head structure, and dynein outer arm composition. No significant differences in expression were observed between aCSF control and IVH/PHH conditions at 72 hours. **D-H**, Graphical representations of normalized counts for select genes involved in neurogenesis and cell proliferation. There was significantly lower expression of Dlx1, Dlx2, Igfbp1, Pax6, and Sp8 72 hours post-IVH/PHH compared to the aCSF control condition. **I-V**, Graphical representations of normalized counts for select genes involved in innate immune activity demonstrates wide upregulation 72 hours post-IVH/PHH compared to the aCSF control condition. For all graphs, adjusted p values are shown with significance = padj < 0.05. Data are shown as mean ± S.D., with n = 3 animals each for the aCSF control and IVH/PHH conditions.

In addition, several key neurogenesis and cell proliferation genes were downregulated in response to IVH-PHH, including *Dlx1 and Dlx2* (p<0.001, Figure 6D), *IgfbpI1* (p<0.001, Figure 6E), *Pax6* (p<0.001, Figure 6F), and *Sp8* (p<0.001, Figure 6G). Of note, *Neurod1*, which has previously been shown to be essential for adult neurogenesis and converting reactive glial cells into functional neurons among other neural transcription factor functions^35–37^, was upregulated after IVH-PHH compared to controls (p=0.026, Figure 6H).

### IVH-PHH does not significantly disrupt cilia-mediated CSF flow domains in the lateral ventricle

While mammalian cilia-based flow networks have been defined in the third ventricle^38,39^, the lateral ventricle, which is the primary site of ventriculomegaly and injury in our model of IVH-PHH^26^, is less well-defined. To identify motile cilia-mediated flow patterns in the lateral ventricle at baseline, we performed bead flow analyses using 4 *µ*m fluorescent microbeads on lateral wall explants in aCSF controls. We identified three major flow domains: rostral, superior, and caudal (Figure 7A). In general, the rostral domain consisted of one or more swirls, in most rodents including at minimum a rostral counterclockwise swirl and/or a clockwise caudal swirl (Figure 7A). The superior domain consisted of a large clockwise swirl (Figure 7A). The caudal domain was more heterogenous, with some animals demonstrating high velocity straight-line flow towards the inferocaudal end of the lateral ventricle (Figure 7B) while other animals had a single large counterclockwise swirl (Figure 7A).

**Figure 7.**
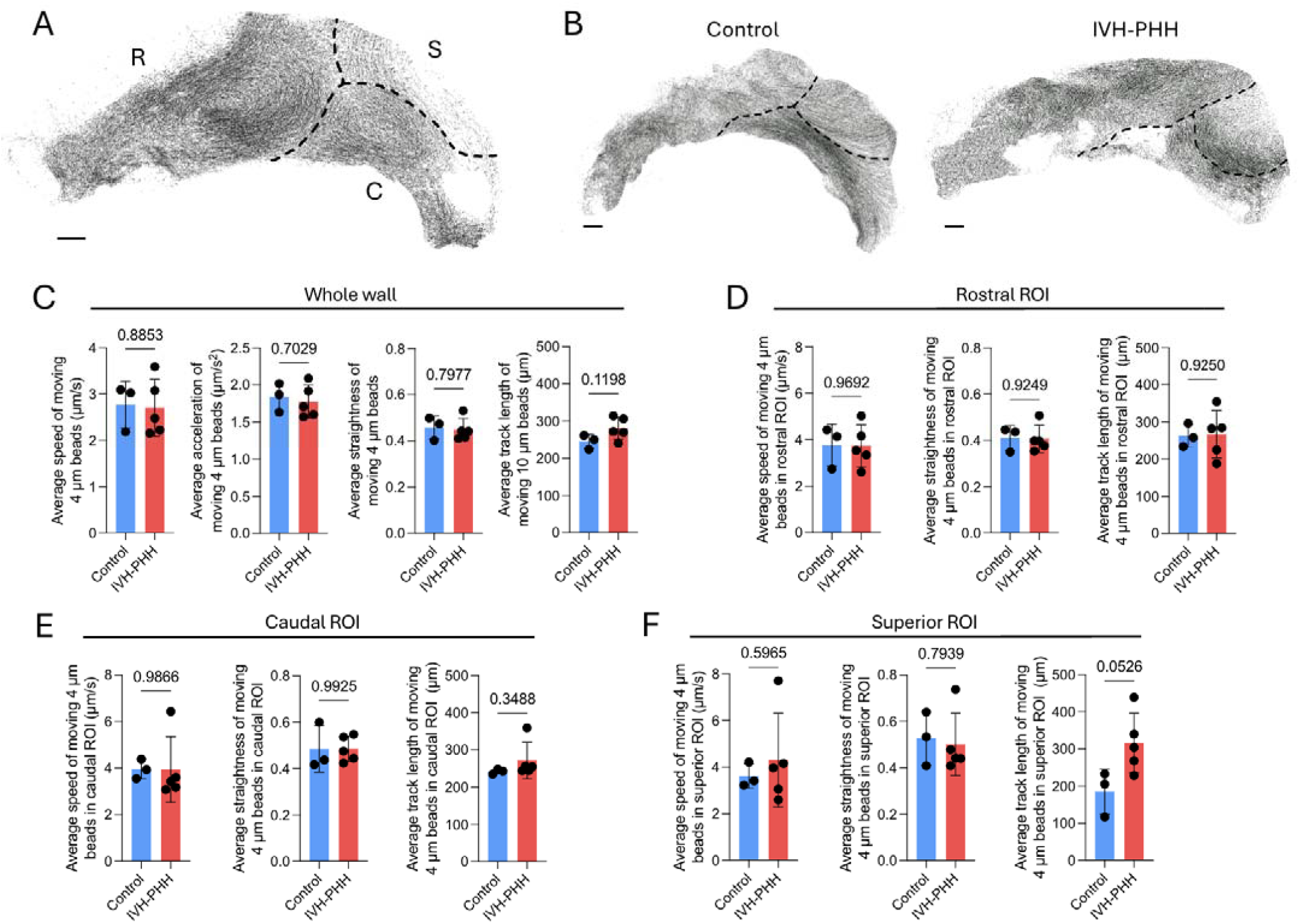
Lateral ventricle lateral wall is divided into rostral, caudal, and superior cilia-mediated flow domains, which are not significantly affected by IVH-PHH. **A**, Flow map demonstrating ex vivo cilia-mediated track displacement of 4 um fluorescent microbeads on the lateral wall of the right lateral ventricle of an aCSF control rat. The lateral ventricle is subdivided into three flow domains, 1. a rostral (R) domain consisting of one or more swirls including a rostral counterclockwise swirl and a caudal clockwise swirl, 2. a superior (S) domain, and 3. a more variable caudal (C) domain consisting of straight-line flow in some animals and a large-radius counterclockwise swirl in others. **B**, Representative flow maps of cilia-mediated displacement of 4 um fluorescent microbeads 72 hours after aCSF control (left) or IVH-PHH induction (right). **C**, Average speed, acceleration, straightness, and track length of 4 um fluorescent microbead cilia-mediated flow across the whole lateral ventricle lateral wall of aCSF control and IVH-PHH rats 72 hours post-induction. **D-F**, Average speed, straightness, and track length of 4 *µ*m fluorescent microbead cilia-mediated flow across the rostral (D), caudal (E), and superior (F) ROIs. All data in C-F are mean +/- SEM, n=3-5 rats per condition. Unpaired, two-tailed t-test.

Next, having established IVH detrimentally affects the number of ependyma with motile cilia on their apical surface without affecting the beating function of the individual remaining cilia, we investigated whether IVH alters the lateral ventricle motile cilia-mediated CSF flow domains we identified. We performed bead flow analyses with 4 *µ*m and 10 *µ*m fluorescent microbeads on lateral wall explants isolated from rats induced with aCSF control and IVH-PHH 10 days after induction. We quantified the average speed (*µ*m/s), acceleration (*µ*m/s^2^), track straightness, and track length (*µ*m) of the moving 4 *µ*m beads across the whole lateral wall of the lateral ventricle and found no differences (Figure 7C). Similarly, we found no differences in the average speed (Supplementary Figure 1A), acceleration (Supplementary Figure 1B), track straightness (Supplementary Figure 1C), track length (Supplementary Figure 1D), or speed variation (Supplementary Figure 1E) of the moving 10 *µ*m beads across the whole lateral wall.

We also investigated if there were differences in cilia-mediated flow within each of the three domains by isolating our analyses to ROIs which included only the domain of interest. Similar to whole-wall data, we found there were no differences in the average speed (*µ*m/s), straightness, or track length (*µ*m) of the 4 *µ*m microbeads in the rostral (Figure 7D), caudal (Figure 7E), nor superior (Figure 7F) domains. There were also no differences when the rostral, superior, and caudal domains were compared against each other for both control and IVH-PHH animals (Supplementary figure 2). Together, this data suggests that IVH does not lead to functional changes in motile cilia-mediated CSF flow domains in the lateral ventricle.

## Discussion

In this study, we used histologic, ultramicroscopic, functional, and transcriptional assays to characterize the effect of IVH-PHH on cilia in the lateral ventricle of neonatal rodents both *in vitro* and *in vivo*. We report that IVH leads to fewer ependymal cells displaying cilia on their apical surface, but that this reduction in cilia on ependymal cells does not lead to functional differences in motile cilia-mediated flow along the lateral ventricle wall. In addition, we show that IVH-PHH leads to reduced Foxj1 protein expression in the lateral wall of the lateral ventricle 72 hours after postnatal day 4 induction, but that there are no corresponding changes in *Foxj1* gene expression or in the expression of other cilia-related genes. Transcriptional changes are instead specific to upregulation of various innate immune-associated markers and downregulation of neurogenesis/cell proliferation genes.

Neonatal IVH is a complication of prematurity and commonly occurs within the first three days of life^40–42^, exposing the ependymal surface to blood breakdown products during the period before and/or during motile ciliogenesis and maturation. PHH develops within 7-14 days of IVH^43,44^. In conjunction with our *in vitro* data showing decreased number of differentiated, multiciliated cells in response to hemoglobin and our *in vivo* data showing fewer Foxj1+ cells in the lateral wall of the lateral ventricle after postnatal day 4 IVH, it is possible that hemoglobin disrupts ependymal cell differentiation leading to impaired ciliogenesis within the timeframe between IVH and PHH onset. An alternative mechanism by which IVH may lead to the decreases in cilia along the lateral ventricle, supported by our bulk RNA sequencing data showing innate immune upregulation, is through hemorrhage product-induced inflammation, immune infiltration, and/or mechanical damage through direct blood contact with the ependyma causing sloughing of the cilia which had already formed at the time of hemorrhage. However, given our functional bead flow analyses demonstrate that there is no difference in motile cilia mediated CSF flow along the lateral wall of the lateral ventricle despite decreases in cilia, it is not likely that either of these potential effects of hemoglobin on cilia play a major role in altering CSF dynamics to underlie PHH development. Important to this conclusion is that our use of the term CSF flow is distinct from CSF dynamics; we refer to CSF flow as fluid movement we definitively know to be motile cilia-mediated and relatively localized to the ventricle wall, while CSF dynamics refers to global CSF circulation that may or may not be driven by motile cilia. All together, we propose that the decreases in cilia are not likely to be a major contributor to global CSF dysfunction and PHH pathogenesis because our experiments demonstrate that cilia-mediated CSF flow is not altered after IVH.

One explanation for these results is that the degree of cilia loss induced by IVH is not sufficient to alter CSF flow. Specifically, through SEM and TEM analyses, we found that while much of the ependymal surface was void of cilia after IVH-PHH, many ciliary tufts consisting of long cilia with preserved ultrastructure remained. Our *in vitro* beat frequency suggest that the function of these remaining cilia is not altered. Therefore, it is possible that the beating action of the remaining cilia can compensate for the lost cilia in maintaining normal cilia-mediated flow along the ventricle wall. Alternatively, this data may suggest that cilia do not play an important role in mediating CSF dynamics in the ventricles. Future bead flow experiments with lateral wall explants isolated from a genetic mouse model where all cilia function is knocked out are needed to gain additional insight on this as a possibility. Finally, while we analyzed the movement characteristics of two different sized beads and found no large differences between control and IVH-PHH conditions, it is possible that we were unable to detect small local changes in flow with the methodology used.

To this point, the contribution of cilia as primary drivers of directional CSF movement has recently been questioned, particularly in the context of brain development and congenital hydrocephalus^18–20^. Cilia are not present throughout fetal development and are not functionally mature in mice until approximately postnatal day 7^18,33^. In addition, hydrocephalus pathology is often present before motile cilia development^19,20,45–51^. Motile cilia and their contributions to fluid dynamics therefore cannot primarily underlie CSF circulation and congenital hydrocephalus pathogenesis in fetal and perinatal development. An alternative mechanism of congenital hydrocephalus pathogenesis that has often been proposed is altered neurodevelopment,^18–20^ specifically that mutations in cilia genes also affect signaling and neural stem cell fate leading to perturbed brain structure and cortical integrity that results in fluid dynamic-independent ventricular enlargement^18,20,52–63^. While this mechanism has primarily been proposed in the context of congenital hydrocephalus, our transcriptional data in this present study showing downregulation of neurogenesis and cell proliferation genes after IVH-PHH suggest that altered brain structure and development may also contribute to ventricular expansion in PHH. However, the increased intracranial pressure and macrocephaly that are clinically seen in PHH suggest that altered fluid dynamics and CSF homeostasis may also play an important role in PHH pathogenesis^17,43,64^. Nonetheless, IVH-induced alterations in parenchymal structure, particularly that of periventricular structures underlying the walls of the lateral ventricles like the hippocampus, cortex, and caudate, represent an important potential mechanism of ventricular expansion in PHH that merits further study. Future studies are needed to elucidate the contribution of altered brain structure vs. CSF dynamics in PHH, in conjunction with alternative mechanisms of PHH including iron-mediated toxicity, oxidative stress, inflammation, complement activation, choroid plexus hypersecretion, and others^17,65^.

In lieu of a role in driving CSF circulation through the ventricular system, recent studies have proposed the function of motile cilia-mediated CSF flow is in establishing CSF compartments within the ventricles^66,67^. Our data showing three major flow domains in the lateral wall of the lateral ventricle support this hypothesis, as the swirls in the rostral, caudal, and superior domains generally corral beads into relatively stereotyped regional swirls, preventing large-scale intermixing between the three flow domains. Given the choroid plexus secretes a diverse array of substances into the CSF, and CSF composition varies by location in the ventricle, it is possible that these flow domains serve to direct important substances (ie. trophic factors, proteins, ions) to ependymal regions overlying cell populations that require CSF-derived factors for proper development in a spatially and temporally dependent fashion^68–73^. Furthermore, as we found IVH does not affect the general organization of the flow domains, it is possible that neurotoxic blood breakdown products secreted into the CSF during hemorrhage are directed towards these regions, differentially affecting the parenchymal structures underlying the swirls we identified. In summary, our data together with recent studies describing cilia-mediated flow patterns promoting CSF compartmentalization in the ventricles suggest the role of cilia in the lateral ventricles may be more targeted towards generating microcirculations of solutes and not facilitating bulk movement of the CSF, a function which may be particularly detrimental to development in the context of IVH and other conditions where neurotoxic substances are released into the CSF.

One limitation to this study is that our investigations of cilia-mediated flow on the lateral ventricle wall were conducted *ex vivo* on lateral wall explants, which may not be representative of CSF dynamics in the ventricle *in vivo* where pulsatility, intracranial pressure, CSF production at the choroid plexus, downstream CSF-interstitial exchange, and other factors may need to be taken into account when qualifying the effect of IVH on CSF flow. However, even if it were the case that the observed decreases in cilia were to alter global, physiological CSF circulation, previous studies have shown that disruptions to cilia beating and changes in CSF flow are not necessarily the initiating factor in hydrocephalus development^74^. Instead, the role of cilia in hydrocephalus may be secondary to altered neurodevelopment, choroid plexus function, or a variety of other mechanisms independent of beat-driven fluid movement. In addition, the lateral wall wholemounts that were used for bulk RNA sequencing included the overlying parenchyma. It is possible that the proportion of ependymal cells relative to other parenchymal cells (neurons, glia) was low, and that IVH-PHH-induced changes in ependymal gene expression were not able to be captured in our analysis as a result. Third, while we hypothesized the relative contributions of 1. impaired ciliogenesis via disrupted ependymal differentiation vs. 2. cilia loss secondary to blood breakdown product exposure, we were unable to definitively show a mechanism by which the decreases in cilia on ependyma occur. Future studies should investigate the specific effect of hemoglobin on ependymal cells and how hemoglobin exposure leads to decreases in the number and density of cilia on the ependymal surface. Furthermore, the timeline over which the decrease in ependymal cilia occurs is not clear. Future studies should repeat our immunohistochemical and electron microscopy investigations at shorter intervals between time of IVH and analysis to understand the temporal nature of the effects of IVH on cilia and ependyma. Finally, future studies should investigate the ependymal surface at longer (>2 weeks) timepoints after IVH. While ependymal cells do not divide after differentiation^33^, cilia have previously been reported to regenerate after deciliation secondary to traumatic brain injury^75^. If cilia loss by blood breakdown exposure is a major contributing mechanism by which the observed decreases in cilia occur, it is possible that the ependymal surface may be capable of recovery in the long term. While cilia loss may not lead to functional changes in CSF dynamics that underlie PHH pathogenesis, it is likely that they play important roles in maintaining homeostasis in the ventricular system. Understanding the mechanism by which they are lost in the setting of IVH may therefore be important towards preventing various neurological sequelae and optimizing outcomes.

## Contributions

S.P. performed experiments, analyzed data, and wrote the manuscript; S.R. performed experiments and analyzed data; L.Y. performed experiments; D.A.G. analyzed bulk RNA sequencing data; I.X. analyzed lateral wall CSF flow data; M.G.B. performed in vitro experiments; L.W.S. analyzed bulk RNA sequencing data and assisted with experiments; G.L.H., S.K., G.M.K., S.P. assisted with experiments and data analysis; D.K.G. analyzed bulk RNA sequencing data; P.K. assisted with electron microscopy experiments; M.J.M. designed lateral wall CSF flow data; P.E. assisted with experiments; D.D.L., P.V.B., A.H., S.L.B. M.R.M. designed experiments and supervised work; J.M.S. designed experiments, supervised the work, and wrote the manuscript.

## Acknowledgements

This work was funded by the National Institutes of Health (R01 NS110793 to J.M.S.), Rudy Schulte Research Institute (J.M.S.), K12 432 Neurosurgeon Research Career Development Program (J.M.S), the Hydrocephalus Association (J.M.S), McDonnell Center for Systems Neuroscience at Washington University in 434 St. Louis (J.M.S.), the Children’s Discovery Institute (J.M.S.), and the Washington University in 435 St. Louis Center for Cellular Imaging also known as WUCCI (J.M.S.).

## Data availability statement

The datasets generated during and/or analysed during the current study are available from the corresponding author on reasonable request.

