## Supplementary figures for "Cilia dysfunction in the lateral ventricles after neonatal intraventricular hemorrhage does not lead to functional changes in cilia-based CSF flow networks"

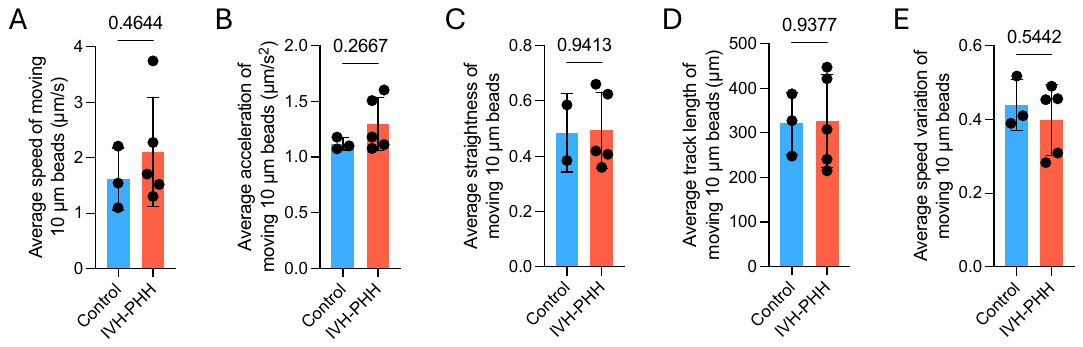

**Supplementary figure 1.** Average speed (µm/s) (A), acceleration (µm/s^2^) (B), straightness (C), track length (µm) (D), and speed variation (E) of 10 um fluorescent microbead cilia-mediated flow across the whole lateral ventricle lateral wall of aCSF control and IVH-PHH rats 72 hours post-induction. All data in are mean +/- SEM, n=3-5 rats per condition. Unpaired, two-tailed t-test.

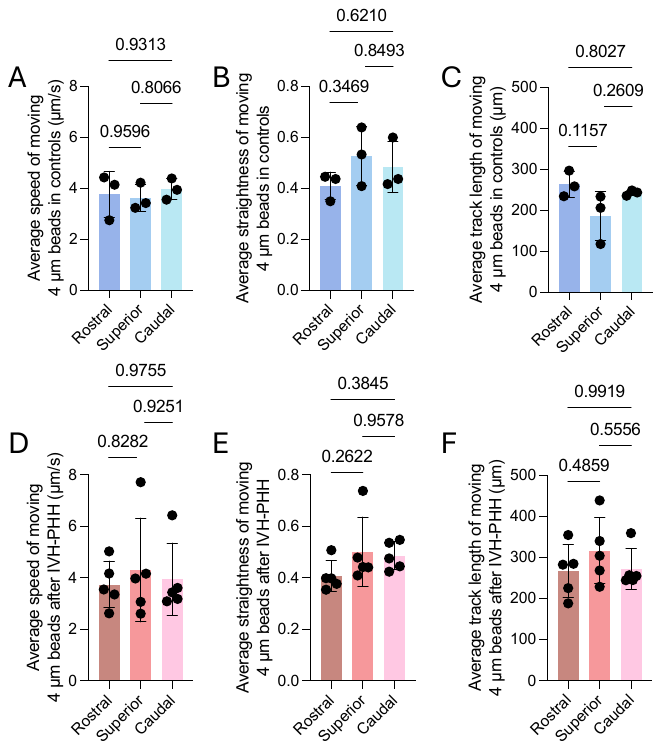

**Supplementary figure 2.** **There are no differences in speed, straightness, or track length between the rostral, superior, or caudal flow domains at baseline and after IVH-PHH. A-C**, Average speed (µm/s) (A), straightness (B), and track length (µm) (C) of 4 um fluorescent microbead cilia-mediated flow in the rostral, superior, and caudal flow domains in aCSF control animals. **D-F**, Average speed (µm/s) (D), straightness (E), and track length (µm) (F) of 4 um fluorescent microbead cilia-mediated flow in the rostral, superior, and caudal flow domains in IVH-PHH animals. All data in are mean +/- SEM, n=3-5 rats per condition. One-way ANOVA with post-hoc Tukey.
